# AgarFix: simple and accessible stabilization of challenging single-particle cryo-EM specimens through cross-linking in a matrix of agar

**DOI:** 10.1101/569087

**Authors:** Klaudia Adamus, Sarah N. Le, Hans Elmlund, Marion Boudes, Dominika Elmlund

**Author notes:** M.B. and D.E. share senior authorship.

## Abstract

Cryogenic electron microscopy (cryo-EM) allows structure determination of macromolecular assemblies that have resisted other structural biology approaches because of their size and heterogeneity. These challenging multi-protein targets are typically susceptible to dissociation and/or denaturation upon cryo-EM grid preparation, and often require cross-linking prior to freezing. Several approaches for gentle on-column or in-tube crosslinking have been developed. On-column cross-linking is not widely applicable because of the poor separation properties of gel filtration techniques. In-tube crosslinking frequently causes sample aggregation and/or precipitation. Gradient-based cross-linking through the GraFix method is more robust, but very time-consuming and necessitates specialised expensive equipment. Furthermore, removal of the glycerol typically involves significant sample loss and may cause destabilization detrimental to the sample quality. Here, we introduce an alternative procedure: AgarFix (Agarose Fixation). The sample is embedded in an agarose matrix that keeps the molecules separated, thus preventing formation of aggregates upon cross-linking. Gentle cross-linking is accomplished by diffusion of the cross-linker into the agarose drop. The sample is recovered by diffusion or electroelution and can readily be used for cryo-EM specimen preparation. AgarFix requires minimal equipment and basic lab experience, making it widely accessible to the cryo-EM community.

**Graphical abstract:** 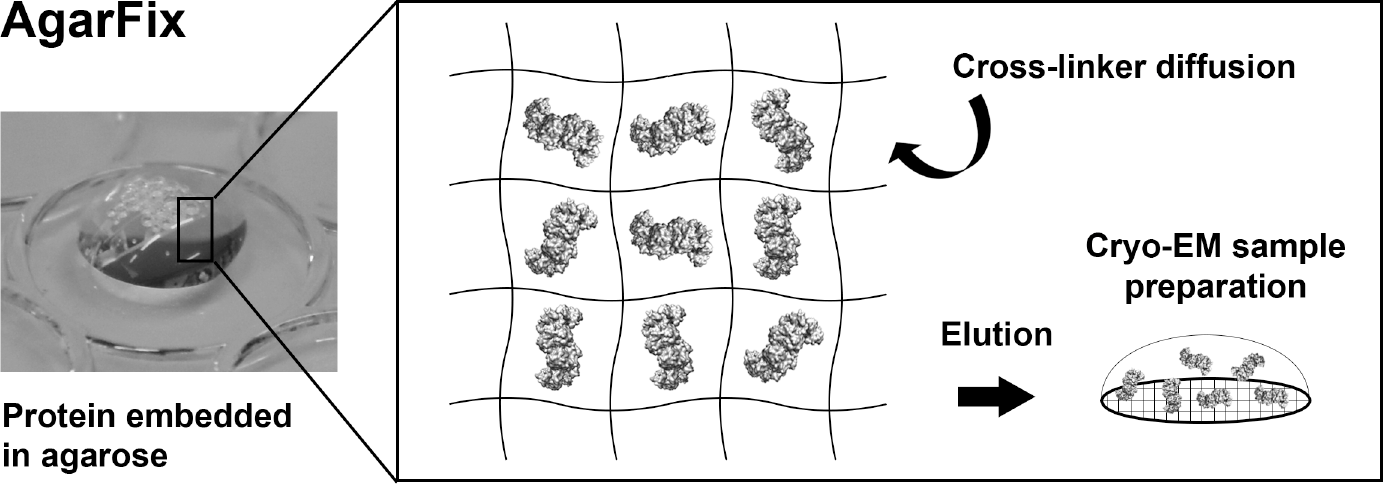

*Highlights:*

- Fragile protein complexes can be cross-linked while embedded in agarose
- The agarose matrix prevents formation of aggregates upon cross-linking
- Cross-linking in agarose is easy, fast, and requires no expensive equipment
- The sample can directly be used for negative stain or cryo-EM grid preparation

## 1. Introduction

Cryogenic electron microscopy (cryo-EM) and single-particle analysis (SPEM) have progressed significantly and is now widely used to generate near-atomic-resolution density maps that allow structure determination at atomic scale of molecules that have resisted other structural biology approaches (Bai et al., 2015; Chen et al., 2018; Liao et al., 2013; Yan et al., 2015; Zhang et al., 2010). However, challenges inherent to complex structural biology targets, such as multi-protein assemblies with dynamic composition and conformation, still remain. These complexes are sensitive to structural damages upon sample freezing, due to adsorption to the air-water interface, causing subunit dissociation and partial to complete denaturation (Taylor and Glaeser, 2008). Fragile samples may also denaturate when exposed to the filter paper during blotting (Palovcak et al., 2018). Avoiding sample degradation during cryo-EM specimen preparation is critical, as it prevents introduction of non-native heterogeneity that would be extremely difficult to distinguish from bona fide functional conformational rearrangements. A common approach to overcome this issue is to use chemical cross-linkers, such as glutaraldehyde (Migneault et al., 2004). Cross-linkers increase the rigidity of complexes by forming covalent bonds between distinct functional groups. However, the direct use of cross-linkers in solution often leads to the formation of unspecific intermolecular cross-links, which can cause aggregations and introduces even greater sample heterogeneity. The GraFix procedure provided a robust solution to this problem by combining gradient centrifugation with chemical fixation (Kastner et al., 2008; Stark, 2010). Briefly, the sample is loaded on top of a glycerol gradient (or other viscous solution, *e.g.* sucrose) containing increasing concentration of cross-linker and subjected to ultracentrifugation. As the sample travels through the gradient, it is exposed to increasing mild cross-linking, while intermolecular cross-linking is avoided due to the centrifugal force pressure (Kastner et al., 2008; Stark, 2010). GraFix is a powerful tool that has successfully been used to cross-link numerous complexes, as illustrated by over 165 citations to date. However, the use of a gradient for cross-linking also presents significant drawbacks. In particular, the high concentrations of glycerol or other viscous agents used in GraFix are not compatible with cryo-EM as they adversely affect the contrast between the macromolecules of interest and the solvent (Drulyte et al., 2018). Therefore, it is essential to remove excess glycerol through dialysis or quick buffer exchange using a desalting column. In our experience, this removal step can detrimentally affect fragile complexes, cause significant sample loss due to precipitation and lead to suboptimal sample quality. Furthermore, reproducible formation of a density gradient and its fractionation require equipment which may be inaccessible to laboratories with limited resources or experimental expertise. Optimizing centrifugation conditions can be tedious, as the sample has to travel just far enough into the gradient to get appropriately cross-linked, but should be distant enough from the bottom to be separated from the aggregates. Consequently, even though very little protein is required to prepare cryo-EM grids, cross-linking using GraFix may require significant investments in time and material. Another alternative is to cross-link the sample while running it through a gel filtration column (Shukla et al., 2014). Briefly, a bolus of cross-linker is injected onto the column and pre-run for a determined volume. The protein sample is then injected so that it runs through the cross-linker. This technique is much faster and might need less optimisation and sample, since an analytical column can be used. However, analytical columns are costly and, in our experience, get damaged and clogged after extensive use for cross-linking. Besides, large complexes cannot always be resolved from aggregates, and the separation properties of gel filtration techniques might dissociate fragile and labile complexes.

We here present an alternative procedure: AgarFix (Agarose Fixation), based on cross-linking of proteins embedded in low-melting agarose. AgarFix takes advantage of the properties of the agarose network, which keeps molecules separated to avoid intermolecular cross-linking, while the cross-linker progressively diffuses into the matrix.

## 2. Materials

### 2.1. Sample embedding into the agarose matrix

Low-melting agarose (*e.g.* SeaPlaque® agarose, Lonza)

Buffer A: cross-linking buffer; compatible with cross-linker (*e.g.* not containing primary amino groups such as TRIS or ammonium sulfate for commonly used aldehyde cross-linkers) 50 mL glass flask or beaker

Heating plate and stirrer, or a microwave

Heat block or water bath

Needle thermometer

96-well tissue culture plate lid: provides an ideal support for handling multiple agarose drops, due to the evaporation rims that prevent adjacent solutions from mixing.

Microcentrifuge tubes

Protein solution at 0.5-2.0 mg/mL

### 2.2. Cross-linking

Cross-linker (*e.g.* glutaraldehyde)

Buffer A’: buffer A supplemented with cross-linker

Quenching agent (*e.g.* Tris, glycine, ammonium sulfate)

### 2.3. Sample recovery

Buffer B: elution buffer; compatible with cryo-EM grid preparation, and, if tolerated by the sample, with lower osmolarity than buffer A in order to increase the elution efficiency.

Scalpel blade

Microcentrifuge

Ultrafiltration centrifugal devices (if sample concentration is required)

### 2.4. Optional step - purification of multi-component complexes prior to cross-linking

Buffer C: buffer suitable for both electrophoresis and protein stability (for example, Tris/glycine native gel buffer)

High gel strength agarose (*e.g.* SeaKem® agarose, Lonza)

Horizontal or vertical gel electrophoresis apparatus

Staining and destaining protein solutions (*e.g.* Coomassie-based)

## 3. Methods

### 3.1. Sample embedding into the agarose matrix

AgarFix requires the use of low-melting agarose, which remains liquid at low temperature (typically above 30°C). For medium to large complexes (> 0.5 MDa) we use SeaPlaque® agarose (Lonza) at a final concentration of 1 %, but for smaller complexes or single proteins, NuSieve® agarose (Lonza) can alternatively be used at 1.5 to 2 % final concentration. The agarose solution is prepared at 1.25X concentration (*i.e.* 1.25 % for a final concentration of 1 %) in buffer A. Agarose melting can be achieved in a microwave or on a hot plate. In either case, the flask containing the solution should be weighted before heating up, and again after melting; if necessary, ultrapure water should be added to bring the flask back to its original weight. The agarose solution is poured into microcentrifuge tubes and kept in a 35°C water bath or heat block until its temperature reaches 37 to 35°C. A needle thermometer can be directly inserted into the agarose solution for precise estimation of the temperature, as it is crucial to avoid overheating the protein sample. The following steps should be performed as quickly as possible to avoid premature agarose gelling: 5 µL of protein solution at 0.5-2.0 mg/mL concentration is pipetted at the centre of the rim of an inverted 96-well tissue culture plate, and 20 µL of melted agarose is added to the protein drop and mixed by pipetting (Fig. 1A).

**Fig. 1.**
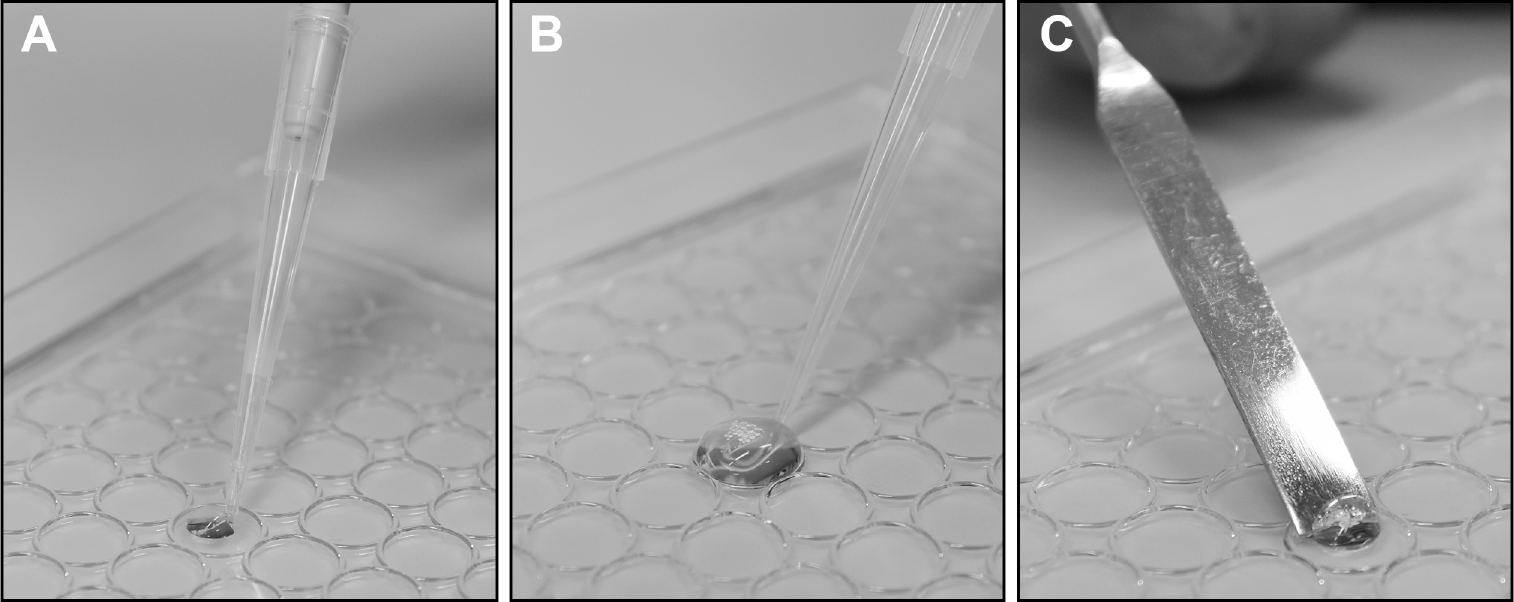
AgarFix procedure. **(A)** 20 µL melted agarose is added to 5 µL protein solution and mixed by pipetting up and down. **(B)** Basic AgarFix step: 125 µL solution (wash, cross-linker, or quenching agent) is added to the agarose drop and, after incubation, is carefully pipetted away. **(C)** Following quenching, the agarose drop is recovered using a spatula.

Since melted agarose cools down to room temperature almost instantly, heating of the protein sample is minimal. Air bubbles may be formed during the mixing process; they are not detrimental and can help locating the agarose drop during the subsequent steps. The amounts stated above are given as a guide; the agarose drop size can be up- or downscaled, and the protein concentration might require optimisation. As a rule of thumb, it is recommended to aim for 0.1-0.5 µM final concentration to avoid the formation of aggregates during the cross-linking process, but this is highly protein-dependent; larger complexes that are prone to aggregation should be kept at low concentrations. Agarose solution aliquots may be stored for a few days at 4°C and reheated in a heat block when required.

### 3.2. Cross-linking

The following steps consist of soaking the agarose drop in buffers A, A’, and B. This can be easily achieved by pipetting buffer onto the agarose drop (Fig. 1B); we found 125 µL to be an ideal volume when using a 96-well plate lid with evaporation rims, but the volume can be adapted depending on the type of the support used. To remove the solution, the pipette tip is placed on the side of the rim and the buffer is pipetted away. Care should be taken not to damage the agarose drop in the process. The lid may be kept on ice, if required for protein stability.

If the protein was initially in a different buffer not compatible with cross-linking (*e.g.* containing primary amines), we recommend pre-washing the agarose drop in 3 × 125 µL buffer A for 3 × 5 min prior to crosslinking.

Cross-linking is achieved by adding to the agarose drop 125 µL of buffer A supplemented with the desired cross-linker; we typically use 0.1% glutaraldehyde. The cross-linker progressively diffuses into the agarose drop, providing gentle fixation. Complete cross-linking is generally achieved within 5 min. When optimising the cross-linking conditions, cross-linking efficiency may be verified by adding the agarose drop to Laemmli sample buffer, heat the sample at 95°C for 5 min, and loading it on a SDS-PAGE gel while still melted. The SDS-PAGE gel can be run in standard conditions. The protein sample will elute from the agarose and migrate into the gel. Following gel staining, visualisation of individual subunits is indicative of incomplete cross-linking; in this case, the cross-linking time should be extended or cross-linker concentration increased. However, we do not recommend extending the cross-linking step more than necessary as artefacts may be caused by over-cross-linking the sample.

After the removal of the cross-linking solution, residual cross-linker may be quenched by adding to the agarose drop 125 µL buffer A supplemented with an appropriate quenching agent (e.g. 50 mM Tris, glycine or ammonium sulfate) for 1 min.

### 3.3. Sample recovery

The simplest option to elute the protein from the agarose drop is by diffusion. The agarose drop is removed from the lid using a small spatula (Fig. 1C) and placed into a microcentrifuge tube containing 50-100 µL buffer B. To increase the speed and efficiency of the recovery, we recommend cutting the agarose drop, and pellet the fragments by centrifugation at 5,000 g for 15 sec. 1 to 2 hours incubation time is sufficient for sample elution, but this vastly depends on the sample molecular weight and on the A and B buffer composition. If required, the recovery may be increased by filtering the sample using a centrifugal device such as Nanosep MF Centrifugal Device - 0.2 µm (Pall). Alternatively, the sample may be centrifuged at 13,000 g for 5 min to pellet the agarose fragments, the supernatant transferred to a new microcentrifuge tube, and can optionally be concentrated using ultrafiltration centrifugal devices.

Electroelution may also be used, in a buffer suitable for both electrophoresis and protein stability (buffer C; for example, Tris/glycine native gel buffer). In this case, the agarose drop is placed into a dialysis bag together with 0.2-1 mL buffer C, immersed in cold buffer C in a horizontal electrophoresis gel unit, and run at 3-5 V/cm for 15 min. Noteworthy, the protein of interest might migrate towards the anode or cathode depending on its pI; if the pI is unknown, agarose drop should be centred in the dialysis bag. After electrophoresis, the protein will be in the bag supernatant, which can be recovered and optionally concentrated. Other recovery methods such as gel nebulisers or freezing should not be used, as they are generally not gentle enough for fragile protein complexes.

When cross-linking a sample for the first time, we recommend assessing the absence of aggregates by negative stain EM. Otherwise, the sample can be readily used for cryo-EM grid preparation.

### 3.4. Optional step - purification of multi-component complexes prior to cross-linking

When preparing complexes from two or more binding partners, it is recommended to eliminate unbound partners in order to minimise sample heterogeneity. This can easily be streamlined by adding a gel-purification step prior to cross-linking. To this end, binding partners are mixed and incubated in conditions promoting their binding, preferentially with one partner in excess. Sample is then loaded in duplicate into a 1% SeaPlaque® agarose (Lonza) horizontal or vertical gel and immersed in cold buffer C. Alternatively, 1% SeaKem® agarose (Lonza) may be used to obtain a stronger, easier-to-handle gel. The gel is run, preferentially with cold buffer, in a cold room or with an active cooling unit, at 2-5 V/cm for 2 hours or longer, depending on the protein molecular weight and charge. One lane is excised and stained with Coomassie dye or derivatives. If required, individual bands can be cut off for isolation and determination of their components (for more details, see (Kim, 2011)). Once identified, the band corresponding to the complex of interest is excised from the duplicate lane, using the first lane as a guide. The purified multi-component sample can be cross-linked using the above-described protocol.

## 4. Results and discussion

Negative stain micrographs obtained from protein complexes uncross-linked, cross-linked in solution, with GraFix and with AgarFix are shown in Fig. 2.

**Fig. 2.**
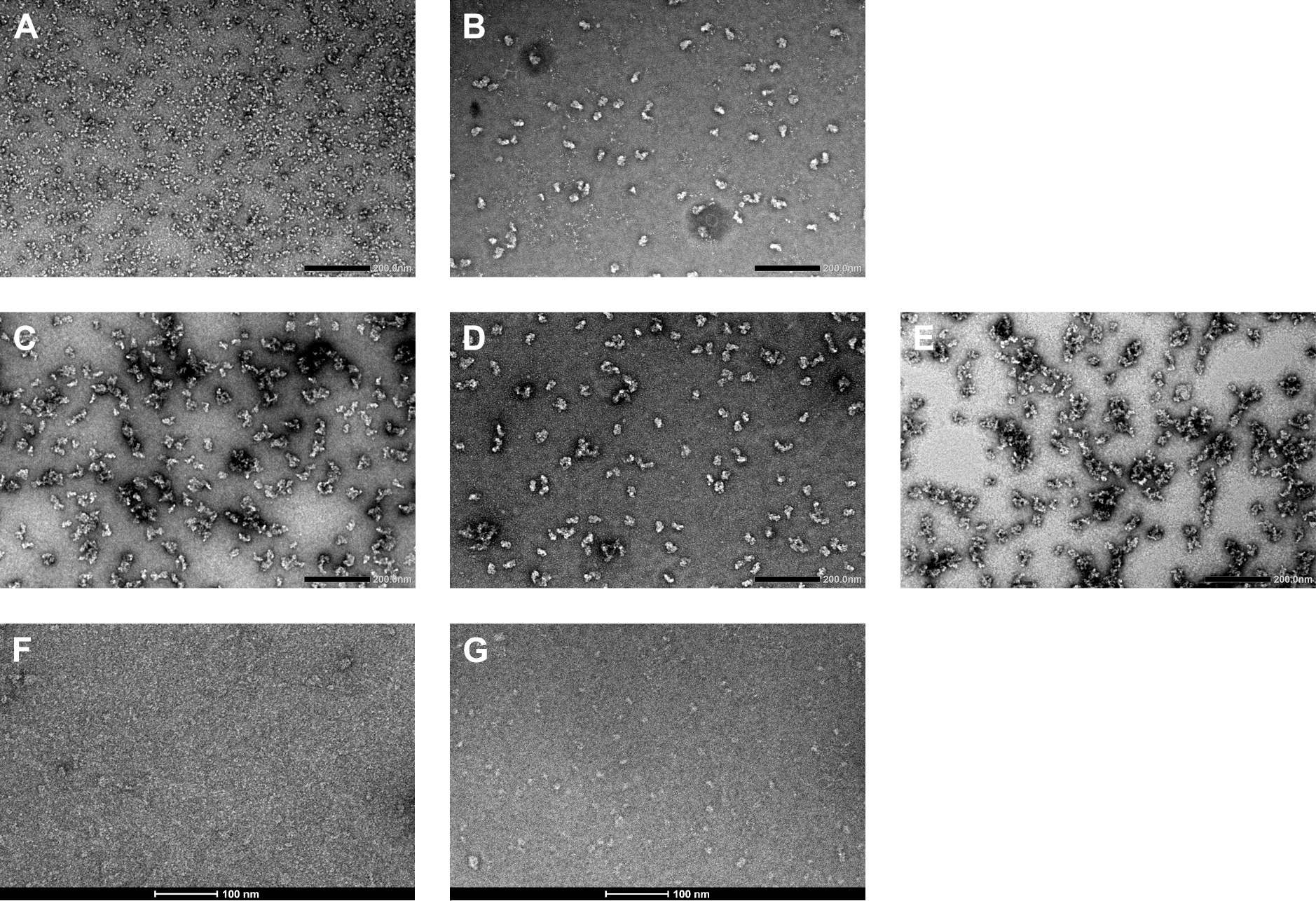
**(A-E) Representative negative stain micrographs of a 1.5 MDa transcription complex (A) uncross-linked** and cross-linked using several approaches. **(B) AgarFix procedure**. 5 µL protein solution at 1 mg/mL was mixed with 20 µL molten agarose (final concentration 0.2 mg/mL), cross-linked in 125 µL buffer A’ for 5 min. Sample was quenched for 1 min in buffer A supplemented with 50 mM ammonium sulfate, and eluted using diffusion into 50 µL buffer A supplemented with 50 mM ammonium sulfate (for consistency purposes between the different approaches) for 60 min and immediately used to prepare negative stain grids. **(C) Cross-linking in solution**. 12.5 µL of buffer A supplemented with 0.2 % glutaraldehyde was added to 12.5 µL of protein at 0.4 mg/mL for 5 min (final glutaraldehyde concentration: 0.1 %; final protein concentration: 0.2 mg/mL). Ammonium sulfate was added to 50 mM for quenching, and sample was immediately used to prepare negative stain grids. **(D) GraFix procedure**. A 10-40 % (w/v) glycerol gradient in buffer A, heavy solution supplemented with 0.3 % glutaraldehyde, was prepared using a gradient former (Gradient Master, BioComp Instruments). 200 µL protein solution at 1.5 mg/mL was loaded onto the gradient and ultracentrifuged in a SW60 rotor for 13 hours at 35 krpm. Gradient was fractionated (Gradient Master, BioComp Instruments) and fractions containing the cross-linked complex (approximately 2/3 into the gradient) were quenched by addition of ammonium sulfate to 50mM and immediately used to prepare negative stain grids. **(E) GraFix procedure followed by glycerol removal**. Sample was processed as for (D), but fractions containing the cross-linked complex were quenched by addition of ammonium sulfate to 50mM, and buffer exchanged to buffer A supplemented with 50 mM ammonium sulfate, using PD Minitrap G-25 columns (GE Healthcare) according to manufacturer’s instructions. After concentration using an Amicon® Ultra 0.5 mL filter, sample was used to prepare negative stain grids. Buffer A is 500 mM potassium acetate, 50 mM HEPES pH 7.6, 10 µM ZnSO_4_. Buffer A’ is buffer A supplemented with 0.1% glutaraldehyde. **(F, G) Representative negative stain micrographs of TFIIE < 200 kDa transcription complex (F) uncross-linked** and cross-linked using **(G) AgarFix procedure**. 5 µL protein solution at 1 mg/mL was mixed with 20 µL molten agarose (final concentration 0.2 mg/mL), cross-linked in 125 µL buffer A’ for 5 min. Sample was quenched for 1 min in buffer A supplemented with 50 mM ammonium sulfate, eluted using diffusion into 50 µL buffer A for 60 min and immediately used to prepare negative stain grids. Buffer A is 150 mM potassium acetate, 50 mM HEPES pH 7.6, 10 µM ZnSO_4_. Buffer A’ is buffer A supplemented with 0.1% glutaraldehyde.

On-column cross-linking could not be included here as this protein complex is too fragile and labile to withstand gel filtration in buffer A or other buffer compatible with aldehyde cross-linkers.

In both cases, cross-linking is strictly required to prevent complex denaturation (Fig. 2A, 2F). Once optimised, cross-linking with AgarFix does not induce aggregates formation (Fig. 2B, 2G), as opposed to cross-linking in solution using the same protein concentration (Fig. 2C). Cross-linking with AgarFix or Grafix results in sample devoid of aggregates, as molecules are kept separated by the agarose matrix or centrifugal force pressure respectively, and gently cross-linked due to the progressive diffusion of cross-linker into the agarose drop or the gradient of cross-linker, respectively (Fig. 2B, 2D, 2G). However, the GraFix method is time-consuming: setting up gradients takes 1 to 2 hours, including solution preparation; gradient are run overnight, following by fractionation and buffer exchange, taking 1 to 3 additional hours depending on the equipment available. In contrast, AgarFix procedure takes 1 to 2 hours. Consequently, cryo-EM grid preparation can occur at the end of the protein purification, rather than the next day. AgarFix is therefore recommended for time-sensitive samples.

A critical difference between AgarFix and GraFix is that AgarFix does not require any dense buffer such as sucrose or glycerol, which are not compatible with cryo-EM grid preparation. In our hands, the glycerol removal step has proven to be problematic for large and fragile complexes and often results in severe aggregation, as shown in Fig. 2E, whether a desalting column or dialysis is used.

Importantly, GraFix is more than a cross-linking procedure, as it also provides an additional purification step. This is critical when preparing samples from two or more binding partners, as unbound components can be purified away as they differentially traverse the gradient according to mass and shape differences. The standard AgarFix procedure does not have this purifying ability; however, multi-component complexes may be gel-purified prior to cross-linking.

### Conclusions

The respective advantages and drawbacks of different cross-linking methods are outlined in Table 1. In summary, AgarFix cross-linking for cryo-EM is a good compromise between the ease of cross-linking in solution and the sample quality achieved with the robust, but more demanding GraFix method.

**Table 1.**
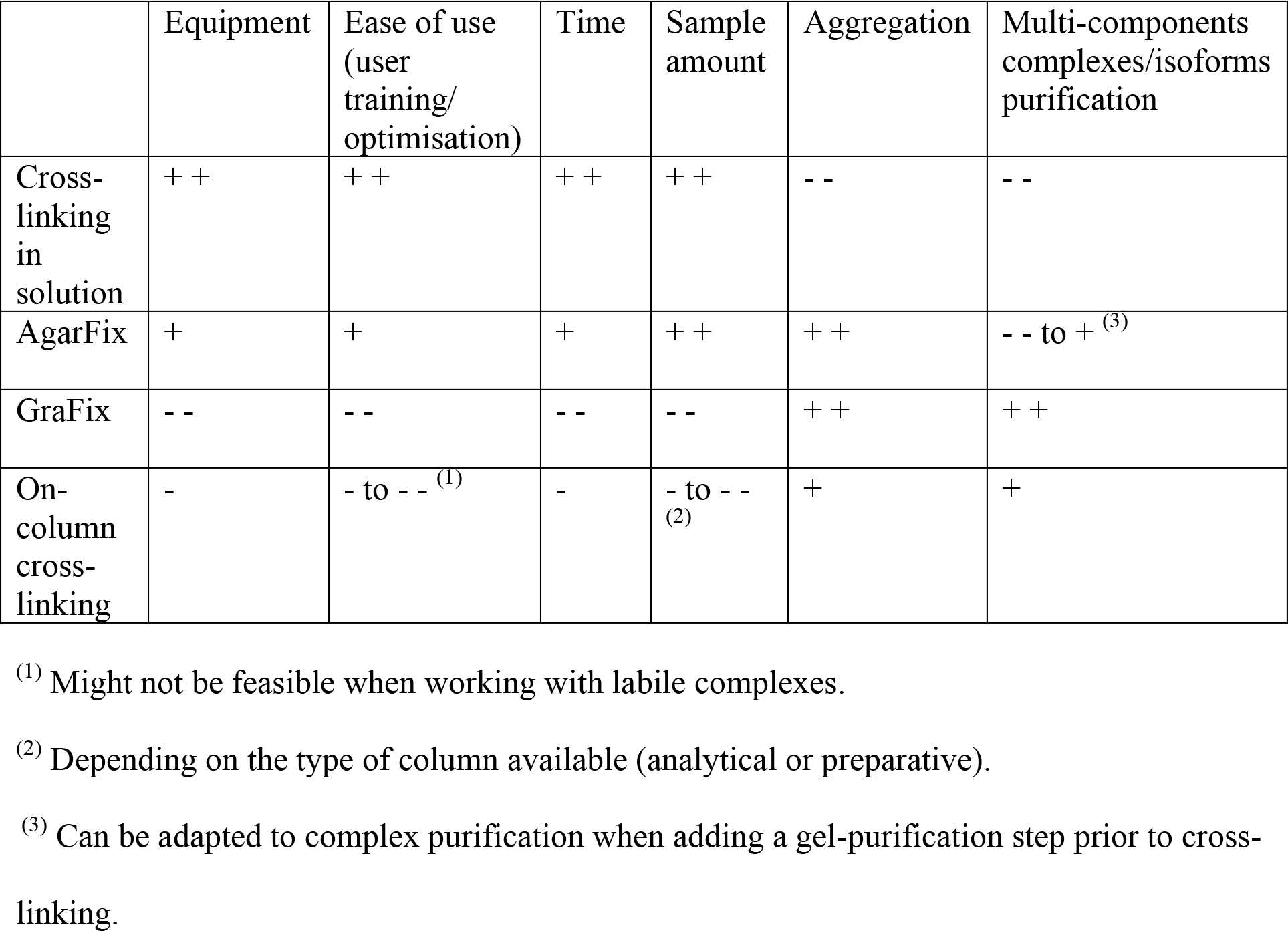
Comparison of respective strengths and weaknesses of cross-linking approaches. Criteria are ranked from - - (most negative) to + + (most positive).

As any other cross-linking method, AgarFix may introduce artefacts and heterogeneity due to differential cross-linking (Hayat, 1986). There are many recent examples of cryo-EM structures that were prepared with cross-linking at near-atomic resolutions (*e.g.* human mTOR complex 2 at 4.9 Å resolution (Chen et al., 2018), human spliceosome at a core resolution of 4.5 Å (Bertram et al., 2017)), some of which could not have been solved without cross-linking (notably RNA polymerase II that dissociates from the uncross-linked pre-initiation complex upon freezing, while cross-linking preserves the complex integrity (Murakami et al., 2013)). We therefore anticipate that easy-to-use and robust cross-linking methods, such as AgarFix, will remain essential in the years to come.

## Acknowledgements

Funding: This work was supported by the Australian Research Council [DE170100701, D.E.; DP170101850, H.E.] and by the National Health and Medical Research Council, Australia [APP1125909, H.E]. M.B. is supported by a Faculty Senior Postdoctoral Fellowship, Monash University, Australia.

